# Sleep and circadian phenotype in people without cone-mediated vision

**DOI:** 10.1101/2020.06.02.129502

**Authors:** Manuel Spitschan, Corrado Garbazza, Susanne Kohl, Christian Cajochen

## Abstract

**Background:** Light exposure entrains the circadian clock through the intrinsically photosensitive retinal ganglion cells, which sense light in addition to the cones and rods. In congenital achromatopsia (ACHM; prevalence 1:30-50,000), the cone system is non-functional, resulting in severe light avoidance and photophobia at daytime light levels. How this condition affects circadian and neuroendocrine responses to light is not known.

**Methods:** In genetically confirmed ACHM patients (n=7; age 30-72 years; 6 women, 1 male), we examined survey-assessed sleep/circadian phenotype (PSQI, ESS, MEQ, MCTQ), self-reported visual function (NEI-VFQ-25), sensitivity to light (VLSQ-8) and use of spectral filters that modify chronic light exposure. In all but one patient, we measured rest-activity cycles using actigraphy over 3 weeks and measured the melatonin phase angle of entrainment using the dim-light melatonin onset (DLMO).

**Results:** ACHM patients experience a severely attenuated light-dark cycle due to severe light sensitivity and habitual use of filters to reduce retinal illumination. In aggregate, both MEQ and MCTQ indicated a tendency to late chronotype. We found regular rest-activity patterns in all patients and normal phase angles of entrainment in participants with a measurable DLMO.

**Conclusions:** Our results reveal that a functional cone system and exposure to daytime light intensities are not necessary for regular behavioural and hormonal entrainment, even when survey-assessed sleep and circadian phenotype indicated a tendency for a late chronotype and sleep problems in our ACHM cohort. Our results can be explained by an adaptation mechanism in circadian photoreception which adjusts to the range of habitual light exposures.

## Introduction

Light exposure at even moderate intensities at night attenuates the production of the hormone melatonin and shifts circadian rhythms in physiology and behaviour (1). Light acts as a *zeitgeber*, enabling entrainment of the circadian clock to the periodic changes in ambient light levels (2). Generally, brighter light has a stronger *zeitgeber* strength, thus providing a more powerful input drive to the circadian timing system (2, 3). These non-visual effects of light on the circadian clock are mediated by the retinohypothalamic pathway, which is largely driven by the intrinsically photosensitive retinal ganglion cells (ipRGCs) expressing the photopigment melanopsin (4). The ipRGCs are ‘non-classical’ photoreceptors signalling environmental light intensity independent of the ‘classical’ retinal photoreceptors, the cones and the rods (Fig. **1*a, b***). The range at which these photoreceptors are active differ (Fig. **2*a***), with cones and ipRGCs responding to moderate to bright light (photopic light levels). Rods, expressing rhodopsin, on the other hand, are 1000 to 10000 times more sensitive and signal in dim and dark light, and saturate at photopic light levels (5).

**Figure 1.**
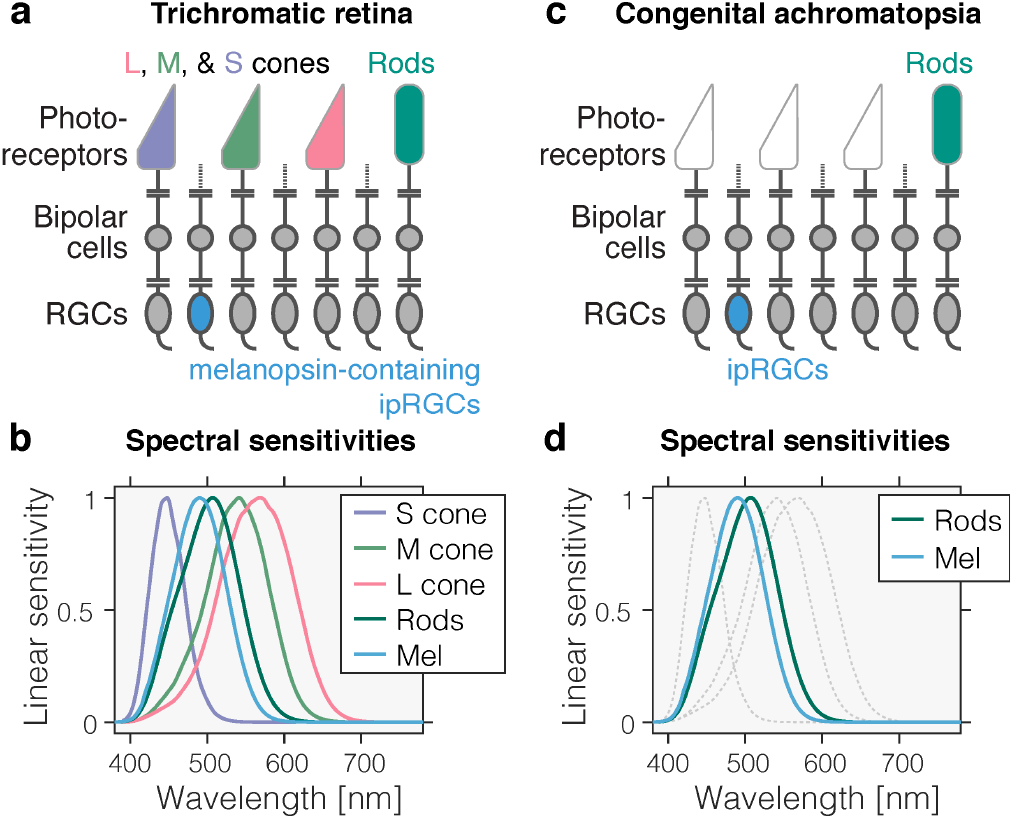
Photoreceptors in the trichromatic and achromatic human retina. ***a*** Schematic diagram of the normal, trichromatic human retina containing three classes of cones – long [L]-, medium [M]-- and short [S]-wavelength-sensitive cones –, rods, and the intrinsically photosensitive retinal ganglion cells (ipRGCs) expressing the photopigment melanopsin. ***b*** Spectral sensitivities of the photoreceptors in the trichromatic retina, showing the overlapping *in vivo* wavelength sensitivity for the S (λ_max_ = 448 nm in linear energy units after pre-receptoral filtering), M (λ_max_ = 541 nm), and L (λ_max_ = 569 nm) cones, the rods (λ_max_ = 507 nm), and melanopsin (λ_max_ = 490 nm). Spectral sensitivities shown here assume a 32-year old observer and include pre-receptoral filtering (52). ***c*** Schematic diagram of the retina of a congenital achromat, missing functional cones, thereby only containing rods and ipRGCs. ***d*** Spectral sensitivities of the photoreceptors in the achromat retina. Faint dashed lines corresponding to the L, M and S spectral sensitivities are given for reference only.

**Figure 2.**
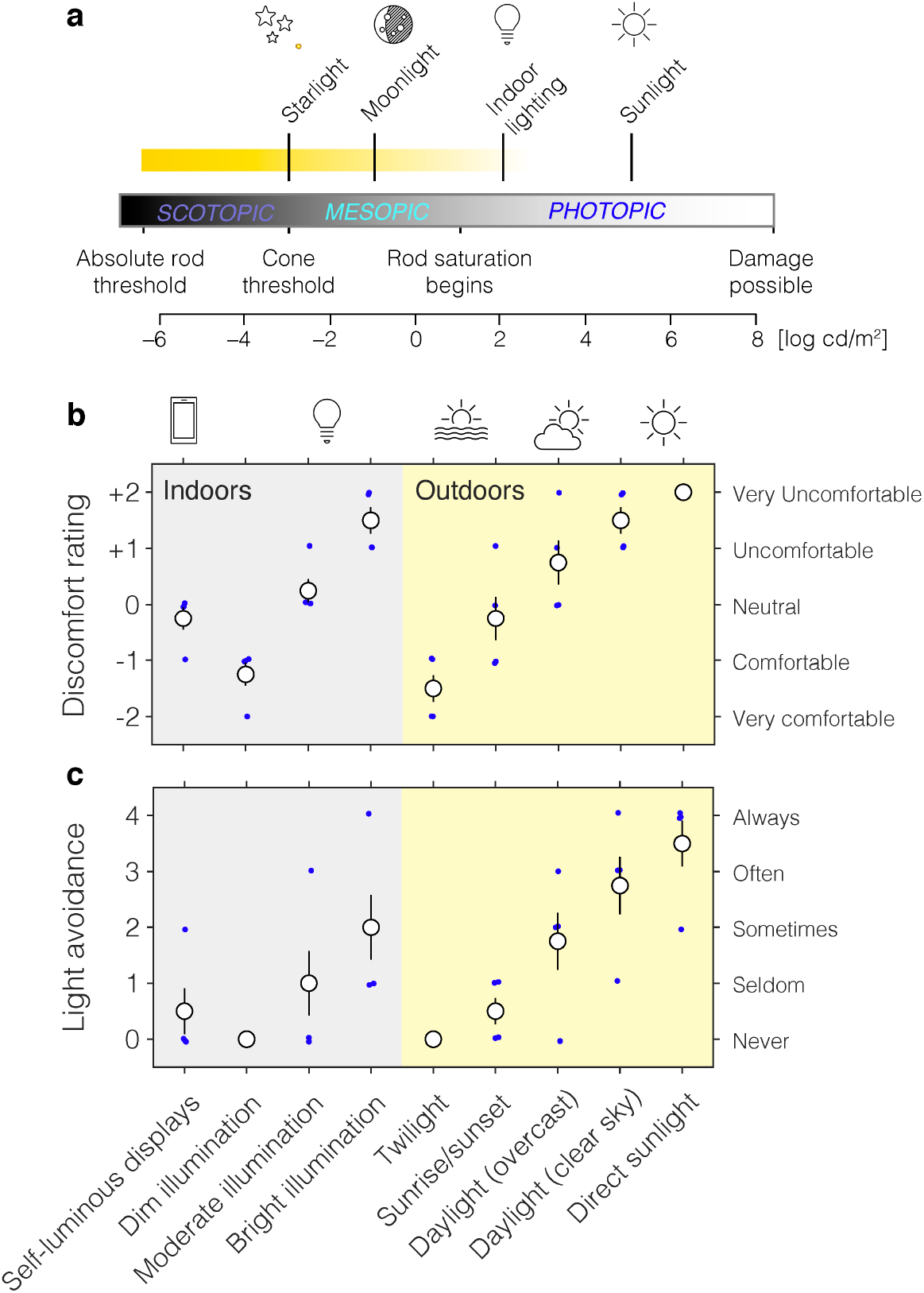
Light sensitivity and light avoidance in congenital ACHM. ***a*** Range of light levels and corresponding environmental conditions. The range of congenital achromats is indicated as a yellow, fading horizontal bar. ***b*** Ratings of light sensitivity and visual discomfort across a range of commonly encountered lighting conditions, indicating severe light sensitivity in bright light. ***c*** Ratings of light avoidance when filters are not used. To manage the hypersensitivity to light, congenital achromats use a range of filters that reduce retinal illumination. Per-participant data on filter use is given in Supplementary Figure **S1**.

The normal trichromatic retina (Fig. **1*a***) contains three classes of cones – the short [S]-, medium [M]- and long [L]-wavelength sensitive cones –, the rods, and the ipRGCs. In *congenital autosomal recessive achromatopsia* (ACHM), also called *rod monochromacy* (estimated prevalence 1 in 30,000-50,000 people (6)), the cone photoreceptors are non- or dysfunctional. In most cases, this is due to mutations in the genes *CNGA3*, *CNGB3*, *GNAT2*, *PDE6H*, and *PDE6C* which affect different aspects of the phototransduction process in cone cells (7). Rarely, mutations in *ATF6* have been shown to also cause ACHM. As the cones are sensitive to moderate to bright lights and responsible for vision of colour, motion and spatial details at daylight light levels, patients with congenital ACHM lack functional photoreception in the upper range of typical day light exposures. This leads to strong visual discomfort, glare and light aversion (8). Congenital achromats are hypersensitive to light (9), with corneal photosensitivity thresholds being 100 to 1000 times lower than for healthy controls (10). To be able to cope with typical, in particular daytime light levels, management of congenital ACHM includes the use of tinted filter glasses (11).

Standard circadian theory predicts that lack of exposure to bright light would reduce the strength of light as *zeitgeber* (2). This prompts the question: If congenital achromats are not exposed to bright light levels, is their circadian photoentrainment impaired? While aspects of rod-mediated visual function in ACHM have been examined before, the question of non-classical photoreception in congenital ACHM has to our knowledge not yet received scientific attention. The authoritative monograph on vision in congenital ACHM does not contain any discussions on non-visual effects of light (8), not least because it predated the discovery of melanopsin. There is, however, anecdotal evidence for an adjustment of the circadian system in congenital achromats: A 1992 *New York Times* article on congenital ACHM stated that “[m]any with the disorder are proud night owls, who love going out after dark” (12), and a publication by *The Achromatopsia Network* suggests that many achromats prefer timing of outdoor and recreational activities to the “magical time of twilight” (13).

Previously, it has been shown that in some individuals who are functionally blind, the melatonin-suppressive effect of light is preserved due to a functioning melanopsin-based ipRGC system even in the absence of cone and rod function (14, 15). Direct evidence for a functional preservation of melanopsin-mediated ipRGC function has also been found in other retinal conditions (e.g. Leber congenital amaurosis, (16)). Importantly, however, these individuals do not necessarily experience the severe discomfort reaction to light typical for ACHM and therefore may indeed be exposed to much more daytime light levels than achromats. We hypothesized that the extreme light sensitivity, light avoidance and ensuing use of filters leads to a reduced light-dark cycle, which translates into a regular but later chronotype. We examined the sleep and circadian phenotype in a group of genetically confirmed congenital achromats (n=7, age range 30-72 years; *CNGB3* [n=5] and *CNGA3* [n=2] genotype), employing a comprehensive suite of self-reporting, actimetry and physiological measurements to arrive at the first picture of how sleep and circadian rhythms are affected by a lack of daylight vision.

## Materials and Methods

### Participants

We recruited participants through advertisements targeted to ACHM patients via the Achromatopsie Selbsthilfeverein e.V., a self-help organisation of achromats, and Retina Suisse. A total of ten patients responded to our adverts and agreed to participate. Of these, nine patients completed the surveys, six completed the observational period and five the at-home melatonin assessment. One participant completed the melatonin assessment in the laboratory. Here, we only consider data from the seven participants with genetic confirmation of autosomal recessive ACHM (Table **2**). All participants underwent remote psychiatric examination by the study physician using the telephone-administered MINI-DIPS-OA (17), none revealing clinical psychiatric problems at the time of test. One participant habitually used trimipramine, which is known to affect sleep but has no known effects on the circadian system.

### Saliva and melatonin assays

Saliva samples were collected at home (n=5) and in the laboratory (n=1) using Sarstedt salivettes (Sarstedt AG, Sevelen, Switzerland). Following the *Sleep Check* protocol (Bühlmann Laboratories AG, Allschwil, Switzerland), participants received written instructions to avoid exposure to bright light (dim lighting from a reading light an television was allowed), to not eat during the collection period and not eat bananas and chocolate in the day of collection, to not consume drinks containing artificial colorants, caffeine (e.g. coffee, black, green and ice tea, cola) and alcohol, and avoid intake of medications containing aspirin or ibuprofen. Participants were instructed to rinse their mouths 15 minutes prior to sample collection, leave the salivette swab in their mouths for 3-5 minutes, and not handle the swab with their hands. Upon extraction of the swab, they were instructed to refrigerate the samples immediately, and ship them using express shipping methods. After arrival in the laboratory, the samples were centrifuged and frozen at −20°. These were then either transferred for analysis to the local laboratory or shipped on dry ice to Groningen (Chrono@Work, Groningen, Netherlands), for determination of melatonin concentrations using a direct double-antibody radioimmunoassay (RK-DSM 2 RIA; Bühlmann Laboratories AG, Allschwil, Switzerland), with detection limit (LoB) 0.3±0.21 pg/ml (n=13). The intra-assay coefficients of variation were 10.1% for at 2.5±0.2 pg/ml (n=15) and 13.3% at 23.4±3.1 pg/ml (n=15). The inter-assay coefficients of variation were 15.4% at 2.4±0.4 pg/ml (n=15) and 10.6% at 24.1±2.5 pg/ml (n=15). The evening melatonin profile was fitted with a piecewise linear-parabolic function using the interactive computer-based hockey-stick algorithm to calculate the individual melatonin onset (v2.4) (18).

### Genetic confirmation

All participants included in the final analysis were genetically confirmed achromats (Table **2**). Five of these were *CNGB3*-associated ACHM patients, while two of them carried mutations in the *CNGA3* gene. Of the six participants who participated in the observational study and the melatonin assessment, five were *CNGB3*-ACHM patients and one was a *CNGA3*-ACHM patient. Genetic confirmation in a research setting was performed by the Institute for Ophthalmic Research, Centre for Ophthalmology, University of Tübingen, Germany.

### Surveys

All survey data were collected and managed using REDCap electronic data capture tools hosted at the University of Basel. Patients completed the Sleep Quality Index (PSQI) (19), the Epworth Sleepiness Scale (ESS) (20), the Morningness-Eveningness Questionnaire (MEQ) (21), the Munich Chronotype Questionnaire (MCTQ) (22), the NEI Visual Function Questionnaire (25 items, NEI-VFQ-25) (23), and the Visual Light Sensitivity Questionnaire-8 (VLSQ-8) (24).

Participants also completed custom visual discomfort and light sensitivity, light avoidance and filter use questionnaires. All three questionnaires used commonly encountered lighting conditions and asked for ratings of visual discomfort without filters, light avoidance without filters, as well as frequency of filter use under these conditions using a 5-item Likert scale. The lighting conditions included were direct sunlight, daylight (clear sky without direct sunlight), daylight (cloudy), sunrise and sunset, and twilight (outdoor category), bright, moderate and dim indoor illumination (indoor category), and smartphones, TV and computer use. Two participants completed this questionnaire over the telephone.

### Actigraphy and sleep diary

Participants wore a Condor ActTrust (Condor, São Paolo, Brasil) actiwatch over the course of the 21-day observational protocol. We restricted our analysis to the time period from 12:00 (midnight) on Day 2 to 12:00 (midnight) on Day 20. We analysed actimetry data reported in the normalized Proportional Integration Mode (PIMn) as follows: We estimated the periodicity of the actimetry data using the Lomb-Scargle periodogram using MATLAB’s plomb function (Mathworks, Natick, MA). Furthermore, to visualise the periodicity (Fig. 4), we fit a sum-of-sinusoids to the time bin-averaged (30-minute bins) and *z* scored data with non-linear least squares using MATLAB’s Curve Fitting Toolbox. We incorporated the fundamental frequency (corresponding to a period length of 24 hours) and the second harmonic (corresponding to a period length of 12 hours). To address nonstationarities in the rhythm which would be masked by bin-averaging and not captured by the Lomb-Scargle periodogram, we also performed a wavelet analysis (25, 26) on the activity data. Additionally, we implemented standard non-parametric analyses of actigraphy-derived activity cycles (27) using the pyActigraphy package (28), calculating intra-daily stability (IS) and intra-daily variability (IV). In addition to wearing an actiwatch, participants completed paper-and-pen sleep diaries during the 21-day protocol, asking for self-reported sleep time and wake-up time.

**Figure 3.**
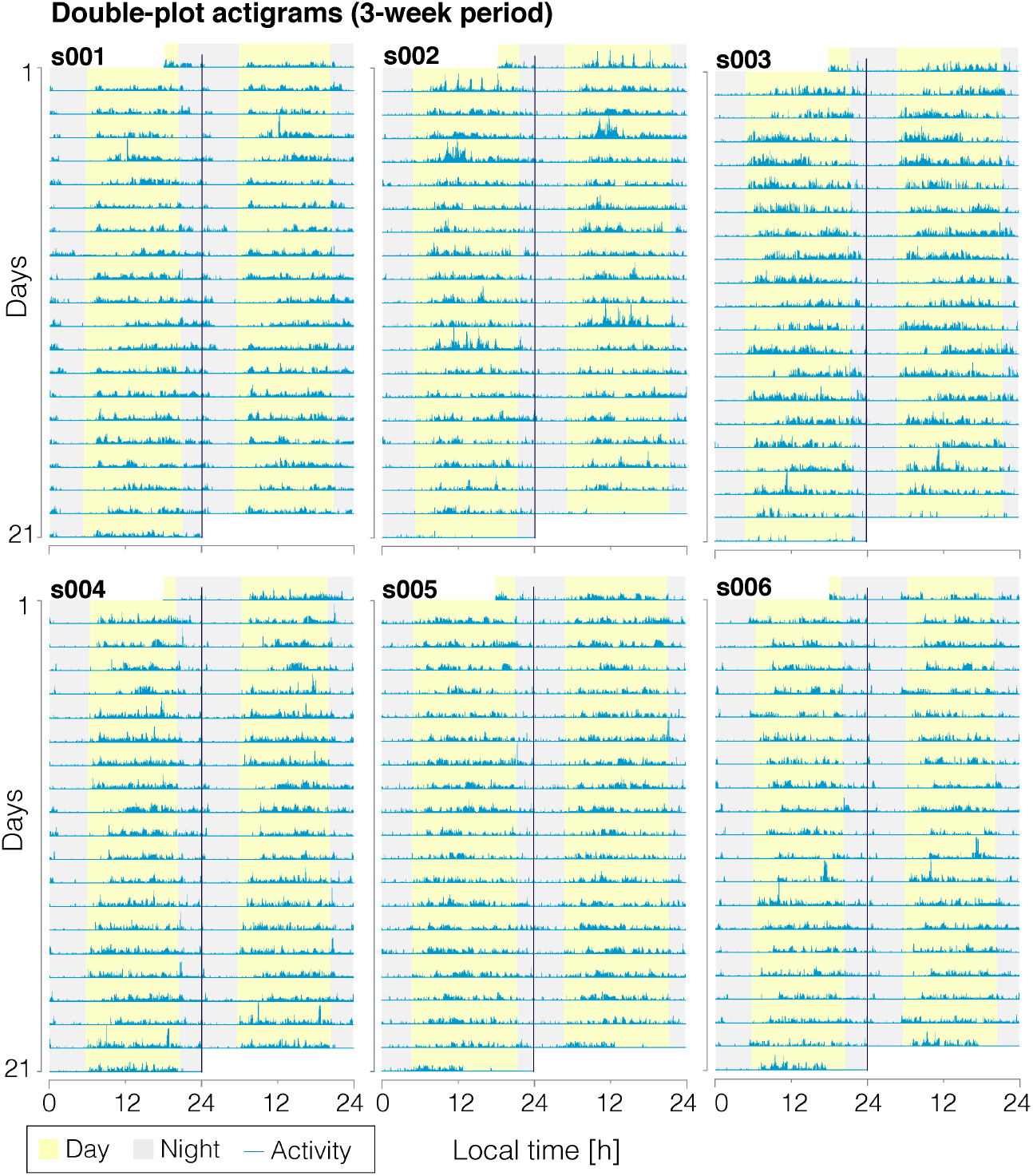
Wrist-referenced actigraphy shows regularity in activity in a group of six achromats (n=6). Data shown across the three weeks of observation. Participants were not instructed to follow a particular rest-activity pattern. Actigrams are shown as double-plots with the x-axis spanning a period of two consecutive days. Shading for day and night is taken from sunrise and sunset times at these chronological dates.

**Figure 4.**
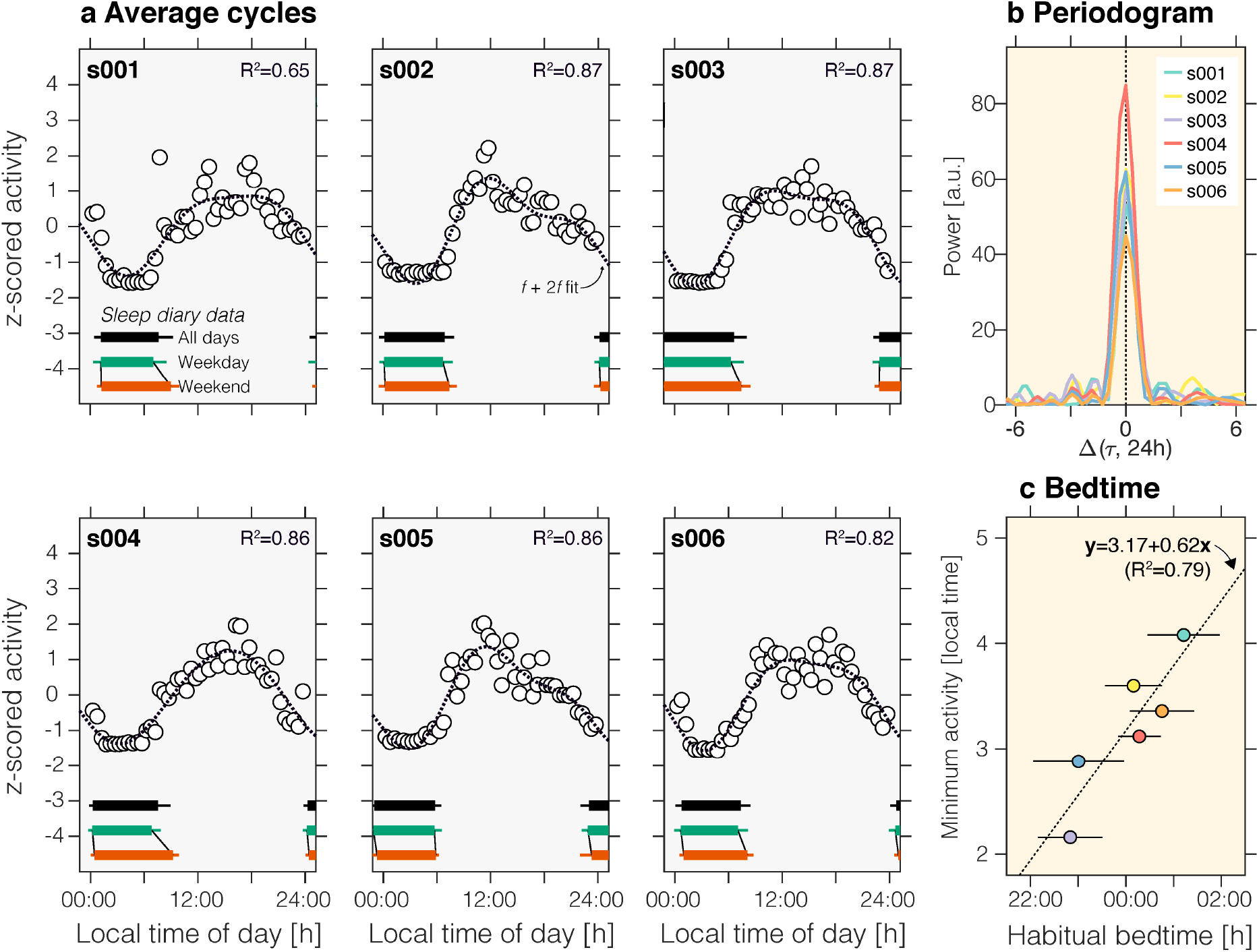
Actigraphy-derived analyses. ***a*** Data from the 21-day observational period were collapsed across days within time of day to yield the average time-of-day activity curves (30 minute bins). For visualisation, averaged data were fitted using a sine-cosine f+2f fit, where f=1/(24.0 hours). All participants show a strong diurnal activity rhythm which can be characterised by a sinusoidal fit on the averaged values (range of R2 values: 0.65-0.87). *Insets*: Average (mean±1 SD, horizontal error bars shown on one side only) bed and wake-up times across the 21-day observational period across all days (black), or aggregated by weekday (green; Monday–Friday), and by weekend (red; Saturday and Sunday). ***b*** Periodogram analysis of actigraphy data, showing a 24-hour period, confirmed by a wavelet analysis in Figure **S4**). ***c*** Relationship between habitual bed time as well as the actigraphy-derived minimum activity timing.

### Ethical approval

Ethical approval for this study was granted from the Ethikkommision Nordwest- und Zentralschweiz (EKNZ), no. 2018-02335. Genotyping in a research setting was approved by the ethics committee of the University of Tübingen, no. 116/2015BO2.

## Results

### Congenital achromats experience an altered light-dark cycle due to extreme sensitivity to light

We confirmed elevated light sensitivity in the sample of congenital achromats, finding high sensitivity to bright lighting conditions, such as direct sunlight, daylight under a clear sky, as well as bright indoor illumination (Fig. **2*a***). This sensitivity to light translates to higher degrees of avoidance of exposure to bright light (Fig. **2*b***). In one patient (s006), we confirmed light sensitivity and retained pupil responses to light in an in-laboratory protocol (Figure **S3**). Together with the survey, this confirms the findings of previous studies, which have found much lower light discomfort thresholds for congenital achromats than for health controls (~3 vs. ~1500 lux) (10) in our sample of patients.

This sensitivity to light is typically managed by the use of optical filters integrated in spectacle glasses or contact lenses. These filters reduce the activation of rods and thereby alleviate visual discomfort and prevent saturation of the rods (29). In Germany, where six of seven of our patients were residing, filters with a transmittance ≤75% and long-pass cut-off filters (cut-off wavelength >500 nm) are prescribable by federal regulation (30) and therefore can be reimbursed through health insurance. In practice, many congenital achromats have at least two filter glasses, a cut-off filter (such as a Zeiss F540, 50% absorption at 540 nm) for indoor use and a cut-off filter with an additional tint (such as Zeiss F90, 90% absorption at 600 nm). In our sample of congenital achromats, we characterised the habitual use of filters using a questionnaire (Fig. **S1**). All participants used a very strong filter to reduce retinal illumination in bright outdoors conditions (Fig. **S1**). Some of our patients (s003 and s005) use up to five separate filters under different conditions, highlighting the complex requirements for management of congenital ACHM, as well as individual differences in light sensitivity.

Due to the substantial overlap of rods and ipRGCs in their response to different wavelengths (Fig. 1***d***), we hypothesized that filter use to reduce rod activation would also reduce ipRGC activation. We tested this hypothesis by examining how spectral filters prescribed in congenital ACHM change the signals of rods and ipRGCs (Fig. **S2*b***). As predicted from the strong overlap and correlation of rod and melanopsin spectral sensitivities, we confirm that rod and ipRGC signals are strongly correlated in everyday light exposures (Fig. **S2*a***). We examined the change of rod and ipRGC signals by simulating the world seen through two common spectral filters (F540 and F90). We find that these two filters reduce the activation of rod and ipRGC signals by a factor of ~0.1× (F540) and ~0.01× (F90) on average, respectively (Fig. **S2*c***). In sum, the use of filters to manage severe visual discomfort in congenital ACHM leads to a significant reduction in habitual light exposure and therefore a significant change in chronic ipRGC activation, the photic driver of the circadian system.

### Survey-estimated chronotype and sleep

We asked participants to complete the Pittsburgh Sleep Quality Index (PSQI) (19), the Epworth Sleepiness Scale (ESS) (20), the Morningness-Eveningness Questionnaire (MEQ) (21), the Munich Chronotype Questionnaire (MCTQ) (22), the NEI Visual Function Questionnaire (25 items, NEI-VFQ-25) (23), and the Visual Light Sensitivity Questionnaire-8 (VLSQ-8) (24). All results are listed in Table **1**. Unexpectedly, we found low scores on the NEI-VFQ-25 (composite score median±IQR [interquartile range] 33.15±4), indicating low vision-related Quality of Life, and high sensitivity to light on the VLSQ-8 (median±IQR: 27±4). For the PSQI, assessing sleep quality over the previous four weeks, we found a range of 3 to 12 (median±IQR: 7±3.5) with 6 above usual cut-off of 5, indicating low sleep-quality. However, only one participant was found to have excessive daytime sleepiness according to the ESS (median±IQR: 6±3). We found a range of MEQ values between 24 and 49 (median±IQR: 40±13), with one definitive evening type, two moderate evening types, and three neutral types. Using the MCTQ, we found a median±IQR mid-sleep MSFsc (mid-sleep on free days) on ~4:00±0.55, corresponding to intermediate/slightly late chronotypes (31). In aggregate, the survey instruments indicate a slight nominal tendency to late chronotypes.

**Table 1:** Demographic details, survey results and participation in sub-studies.

|  |  |  | Sleep |  | Chronotype |  | Visual function |  | Participation |  |  |  |
| --- | --- | --- | --- | --- | --- | --- | --- | --- | --- | --- | --- | --- |
| Patient | Age | Sex | PSQI<br>(19) | ESS<br>(20) | MEQ<br>(21) | MSF<sub>sc</sub><br>(22) | NEI-VFQ-25<br>(Composite<br>Score) (23) | VLSQ-8 total<br>(24) | Filter<br>use<br>survey | Sleep and<br>chronotype<br>survey | At-home<br>DLMO<br>assessment | Laboratory<br>DLMO<br>assessment |
| *s001* | 32 | F | 9 | 6 | 44 | 4.50 | 33.15 | 31 | ✓ | ✓ | ✓ |  |
| *s002* | 64 | F | 3 | 2 | 40 | 3.99 | 32.83 | 19 | ✓ | ✓ | ✓ |  |
| *s003* | 41 | F | 12 | 6 | 46 | 3.60 | 36.67 | 27 | ✓ | ✓ | ✓ |  |
| *s004* | 54 | F | 7 | 9 | 49 | 4.09 | 27.00 | 28 | ✓ | ✓ | ✓ |  |
| *s005* | 72 | F | 7 | 8 | 24 | 3.88 | 35.19 | 26 | ✓ | ✓ | ✓ |  |
| *s006* | 66 | M | 5 | 14 | 31 | 5.00 | 33.61 | 26 | ✓ | ✓ |  | ✓ |
| *s007* | 40 | F | 10 | 6 | 33 | 2.00 | 27.96 | 31 |  | ✓ |  |  |
| Median |  |  | 7 | 6 | 40 | 3.99 | 33.15 | 27 | n=6 | n=7 | n=5 | n=1 |
| IQR |  |  | 4 | 3 | 13 | 0.56 | 4.00 | 4 |  |  |  |  |

**Table 2:**
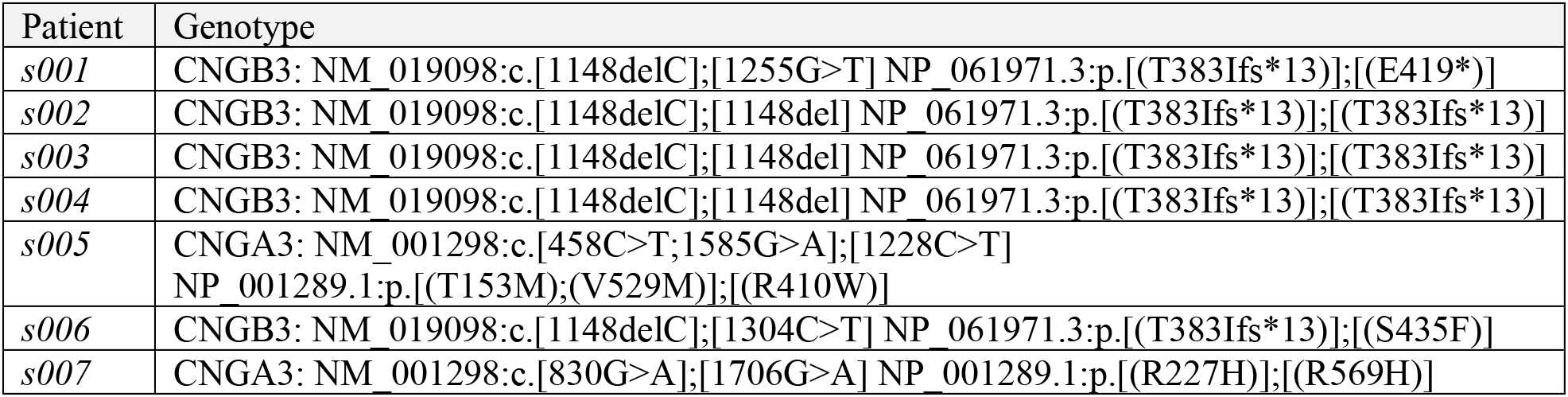
Genotypes of all participants in this study.

| Patient | Genotype |
| --- | --- |
| s001 | CNGB3: NM_019098:c.[1148delC];[1255G>T] NP_061971.3:p.[(T383Ifs*13)];[(E419*)] |
| s002 | CNGB3: NM_019098:c.[1148delC];[1148del] NP_061971.3:p.[(T383Ifs*13)];[(T383Ifs*13)] |
| s003 | CNGB3: NM_019098:c.[1148delC];[1148del] NP_061971.3:p.[(T383Ifs*13)];[(T383Ifs*13)] |
| s004 | CNGB3: NM_019098:c.[1148delC];[1148del] NP_061971.3:p.[(T383Ifs*13)];[(T383Ifs*13)] |
| s005 | CNGA3: NM_001298:c.[458C>T;1585G>A];[1228C>T]<br>NP_001289.1:p.[(T153M);(V529M)];[(R410W)] |
| s006 | CNGB3: NM_019098:c.[1148delC];[1304C>T] NP_061971.3:p.[(T383Ifs*13)];[(S435F)] |
| s007 | CNGA3: NM_001298:c.[830G>A];[1706G>A] NP_001289.1:p.[(R227H)];[(R569H)] |

### Congenital achromats have regular rest-activity cycles

Of our seven participants, six participants completed a three-week long assessment during which they wore actigraphy watches and completed a sleep diary but were not instructed to follow any particular sleep-wake schedule. We found regular rest-activity cycles in all individuals (Fig. **2**). We subjected the actigraphy data to a Lomb-Scargle periodogram analysis (Figs. **4*b***), finding that the rest-activity patterns are periodic with a period length of 24 hours. To examine possible non-stationarities in the rhythm that would not be captured using the periodogram analysis, we confirmed the 24-hour periodicity using a wavelet-based analysis (Figure **S4**). The probability that this 24-hour periodicity is due to participants having a free-running rhythm with 24-hour period is very unlikely (p<0.0001 assuming an exact period of 24h, and p=0.018 for a range of periods in the interval 24±0.2h; Figure **S5**). Additionally, we assessed the regularity and fragmentation of the participants’ activity rhythms (27). The regularity (intra-daily stability; IS: 0.65±0.05 [mean+1SD]) is slightly lower than the estimated population average (32) but significantly higher than a previously characterised sample of psychiatric patients with sleep-wake problems (33). Fragmentation (intra-daily variability; IV: 0.80±0.02) is higher than in the estimated population average (32), confirming the self-reported sleep quality above the normal cut-off.

We further examined whether the sleep-wake rhythms of the congenital achromats exhibit social jetlag (34), which is the phenomenon that participants go to bed and wake up later on the weekends. By drawing on self-reported bed and wake-up times, we found that most of our participants showed on average a delay in their weekend wake-up times (Fig. **4**) of approximately one hour difference in wake-up time on the weekend (median±IQR: 1.12±1.26), but a smaller nominal difference in bed-time (median±IQR: 0.17±0.30).

### Congenital achromats have normal melatonin secretion profiles and phase angles of entrainment

While our actigraphy results strongly point to preserved normal diurnal rhythms in activity and behaviour with a period of 24 hours, these data themselves do not establish that this is due to a preserved circadian rhythm. For example, it is conceivable that the behavioural entrainment can be attributed to non-photic *zeitgebers* (e.g. alarm clocks as a simple example). To rule out this possibility, we examined the secretion of melatonin during the evening hours in a modified at-home DLMO protocol (35). Participants collected saliva every 30 minutes from five hours before to one hour after their habitual bedtime. Samples were refrigerated and shipped to us for biochemical assays (radioimmunoassay for melatonin). In four of six participants who participated in this study, we found a clear rise in melatonin levels (Fig. **4**). The phase angle of entrainment ranged between ~3 hours to ~45 minutes prior to habitual bedtime. This distribution is within the normal range for the melatonin phase angle of entrainment (36). In two participants (s005 and s006; Figure **5**) corresponding to two older patients in our study which habitually take beta-blockers, we failed to detect an increase in melatonin levels in the evening in the expected range relative to habitual bedtime. While this might be due to mistiming of the saliva collection protocol, both individuals had normal rest-activity cycles, suggesting that the lack of measurable DLMO in these participants may be due to the well-known interaction of beta-blockers with melatonin secretion (37) or due reduced melatonin production in elderly participants.

**Figure 5.**
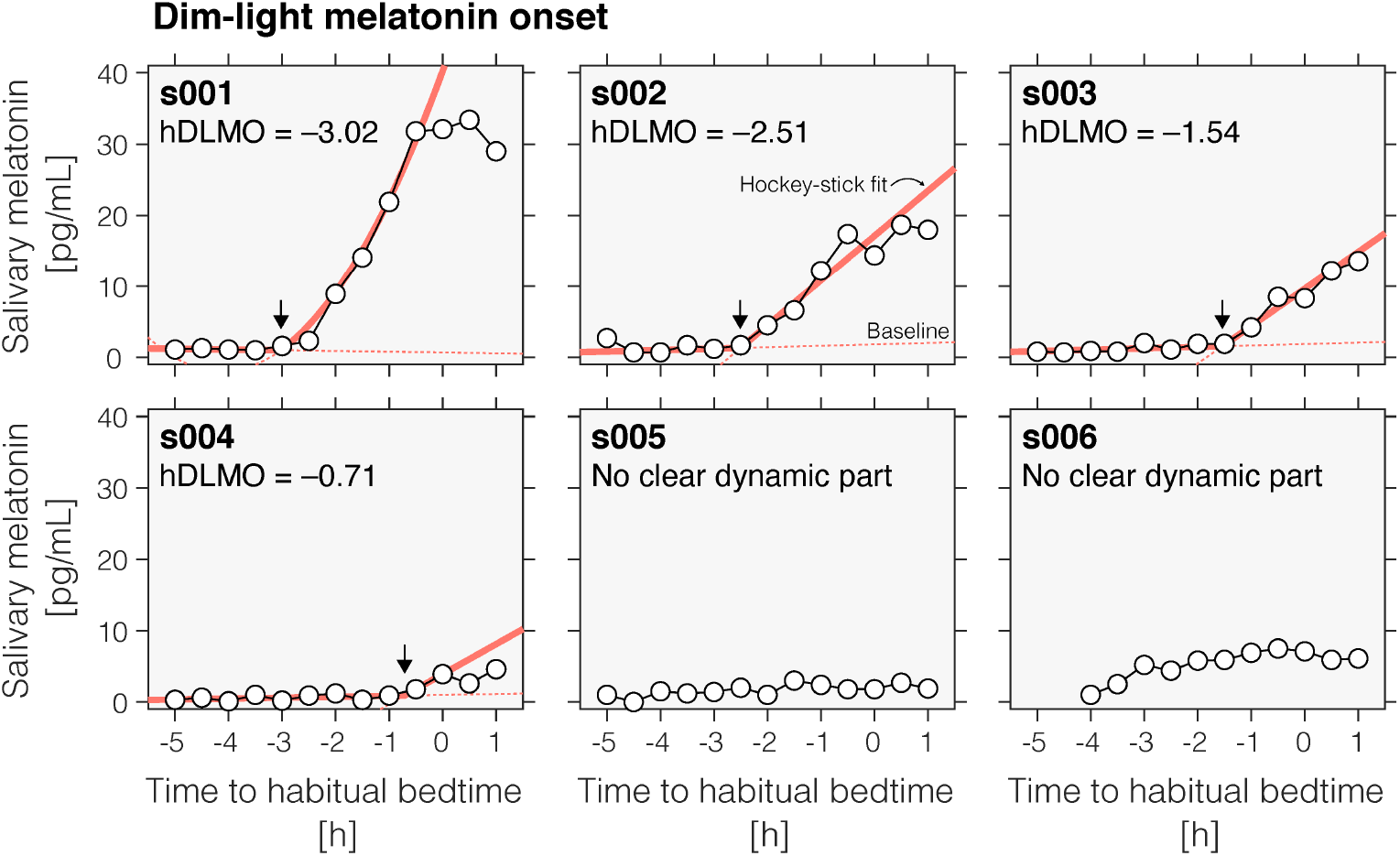
Normal melatonin phase angles of entrainment in four congenital achromats (n=4; total of 6 tested). Dim-light melatonin onset profiles as a function of habitual bedtime, assessed in an at-home measurement protocol using saliva collection. Saliva samples were assayed using radioimmunoassay (see details in text) and DLMO timing was extracted using Hockey Stick software (18). Two of the six participants did not show a clear dynamic rise in their melatonin profiles.

## Discussion

In the first systematic investigation of sleep and circadian phenotype in ACHM, we find that congenital achromats have both normal rest-activity cycles and phase angles of entrainment in the absence of a functional cone system. This adds to our mechanistic understanding of how light fundamentally affects circadian and sleep physiology by demonstrating that low levels of light exposure can support circadian photoentrainment. Importantly, our results show that a functional cone system is not necessary for normally entrained circadian melatonin and sleep-wake cycles in congenital achromats.

In people with a trichromatic retina, entrainment to a 24-hour cycle (but not lower or higher period lengths) can be supported by dim light at around 1.5 lux under very tightly and explicitly controlled conditions (38). Similarly, some laboratory studies performed with pharmacological pupil dilation and dark adaptation have also found very low melatonin suppression thresholds (39). The overwhelming evidence suggests that moderate light exposures are necessary to produce an appreciable effect in circadian entrainment (40, 41) and melatonin suppression. Outdoor light exposure is also systematically related to chronotype (42), with higher outdoor light exposure leading to phase-advanced activity cycles (43). This is consistent with our findings that congenital achromats show a tendency to later chronotypes, likely due to their lack of a strong light exposure signal. Whether rod or cone signals alone are sufficient to influence human circadian and neuroendocrine responses to light in humans is currently not known and will require further investigation. Participants with colour vision deficiencies affecting the L (protanopia) or M (deuteranopia) cones show normal melatonin suppression responses to light, indicating that neither class is necessary for melatonin suppression (44). In animal models, rods have been found to contribute to phase shifting responses (45), thereby effectively extending the range at which light can contribute to circadian photoentrainment.

Behavioural light avoidance and use of filters that reduce retinal illuminance leads to the chronic modification of the “spectral diet” of congenital achromats. A similar adaptation mechanism may tune the sensitivity of circadian photoreception to the range of available light intensities in the environment. In trichromatic observers, chronic modification of retinal input through the use of blue-filtering contact lenses over a two-week period (46) or through the natural ageing (47) leads to an adaptation of the melatonin-suppressive response to light. A similar mechanism is likely at play in congenital ACHM, indicating a flexible gain control mechanism that normalises the sensitivity of the circadian system to the range of habitual retinal illuminances.

The light-dark cycle is the primary driver of circadian entrainment (48), but the circadian system is also sensitive to nonphotic zeitgebers (49), including physical exercise, meal times, and social cues. These, however, may not fully explain our results. It is clear that congenital achromats do experience the light-dark cycle, albeit in an altered way reduced in amplitude. While meal timing can affect the circadian clock (50), these effects may be limited to peripheral oscillators and may not affect DLMO timing (51). The most parsimonious explanation for our results is therefore an adaptation of circadian photoreception to the reduced range of light intensities. Importantly, the results presented here may apply other inherited or progressive retinal disorders and diseases characterised by hypersensitivity to light.

## Acknowledgements

This study was supported by the Wellcome Trust (Sir Henry Wellcome Fellowship to M.S.; Wellcome Trust 204686/Z/16/Z), Linacre College, University of Oxford (Junior Research Fellowship to M.S.), the John Fell OUP Research Fund, University of Oxford (to M.S; 0005460), Retina Suisse, and sciCORE, University of Basel.

We thank Konstantin Danilenko and Evgeniy Verevkin for providing a modified version of their Hockey Stick software, Rafael Lazar for assistance in running in-laboratory studies, Sofia Georgakopoulou from sciCORE, University of Basel for helping with set-up and maintenance of the REDCap survey, Oliver Stefani for photographing the stimulus set up, Forrest Webler for helping with the “spectral diet” measurements, and Roland Steiner for the transmittance measurements. We also wish to thank all volunteers for participating in this research study, and the Achromatopsie Selbsthifeverein e.V. and Retina Suisse for their support.

## Supplementary Appendix

**Supplementary Figure S1.**
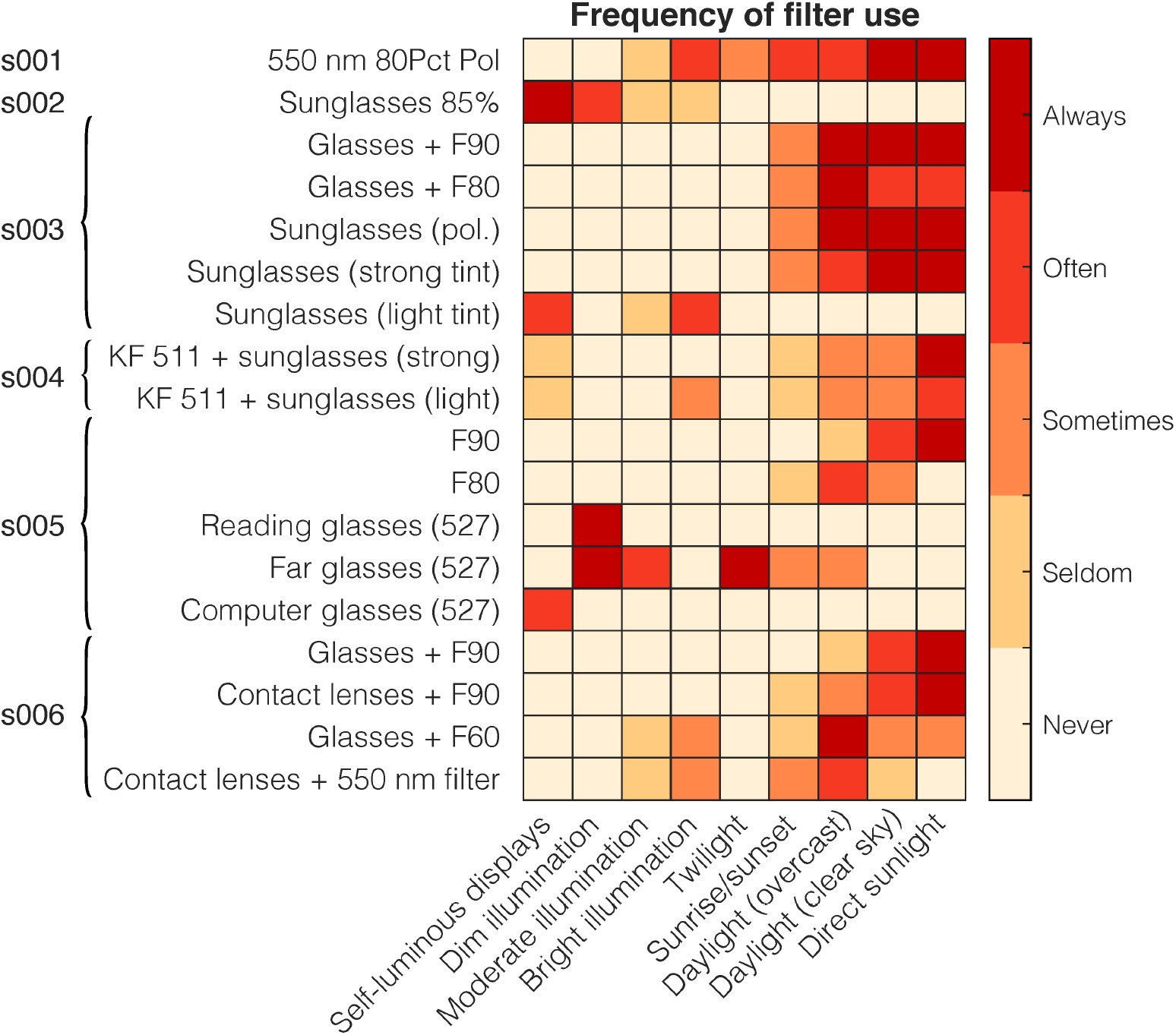
Habitual filter use. Participants were asked to indicate the frequency of filter use under a range of commonly encountered lighting conditions using a 5-item Likert scale. The number of filters used varied widely between participants (1-5 filters used), though all used some filters under daylight intensities.

**Supplementary Figure S2.**
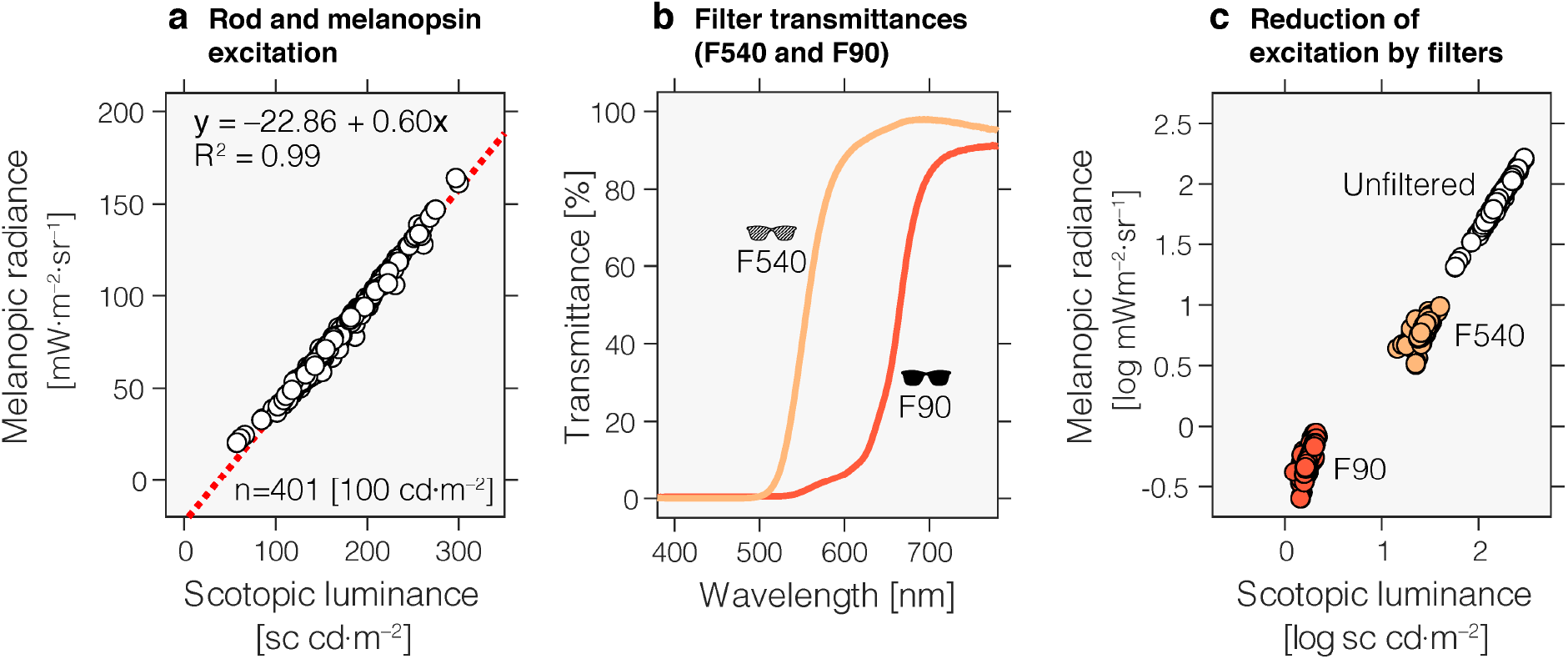
Spectral filters affect rod and melanopsin signals in the achromatic retina. ***a*** Simulation of rod and melanopsin-expressing ipRGC signals under wide range of spectral conditions. We simulated the distribution of rod signals (expressed as scotopic luminance) and ipRGC signals (expressed as melanopic radiance) while pegging the photopic luminance to match 100 cd/m^2^ for 401 spectra representing a wide variety of light sources, including daylight, fluorescent and LED light (53). Under these conditions, rod and melanopsin expressing ipRGC signals are highly correlated and linear with each other, not least owing to the small spectral separation of the rod and ipRGC spectral sensitivities (correlation of their spectral sensitivities: Pearson’s *r*=0.946, p<0.001). Melanopic and scotopic quantities were calculated by matrix-multiplying the spectra with the spectral sensitivities contained in the CIE S 026/E:2018 standard (52). Rod responses were scaled by 1700 lm·W^−1^ (54). ***b*** Transmittances of F540 and F90 tinted filter glasses, which are commonly prescribed in congenital ACHM. The spectral transmittances of the patient-owned F540 and F90 filters were measured between 250 and 2500 nm at 1 nm resolution using a Varian Cary 500 Scan UV-Vis NIR Spectrophotometer (Varian Inc., Palo Alto, CA). ***c*** Simulation of rod and ipRGC signals under the two filters (F90 and F540). The filters reduce both rod and ipRGC radiance by approximately 1 and 2 log units, on average. We confirmed these theoretical calculations using an independent dataset of personalized light exposure measurements with spectral resolution (n=1 healthy control participant; n=4213 usable spectra between 1 and 10,000 lx; clip-on nanoLambda Spectrometer, Daejeon, Korea). For the F540 filter, rod responses are on average reduced by a factor of 0.16× ≈ 0.8 log_10_ units, and melanopic responses are reduced by a factor of 0.08× ≈ 1.1 log_10_ units. On average, The F90 filter reduces rod responses by a factor of 0.01× ≈ 1.98 log_10_ units, and melanopic responses are reduced by a factor of 0.007× ≈ 2.16 log_10_ units, confirming that everyday light exposure profiles yield a reduced melanopic signal with filter usage.

**Supplementary Figure S3.**
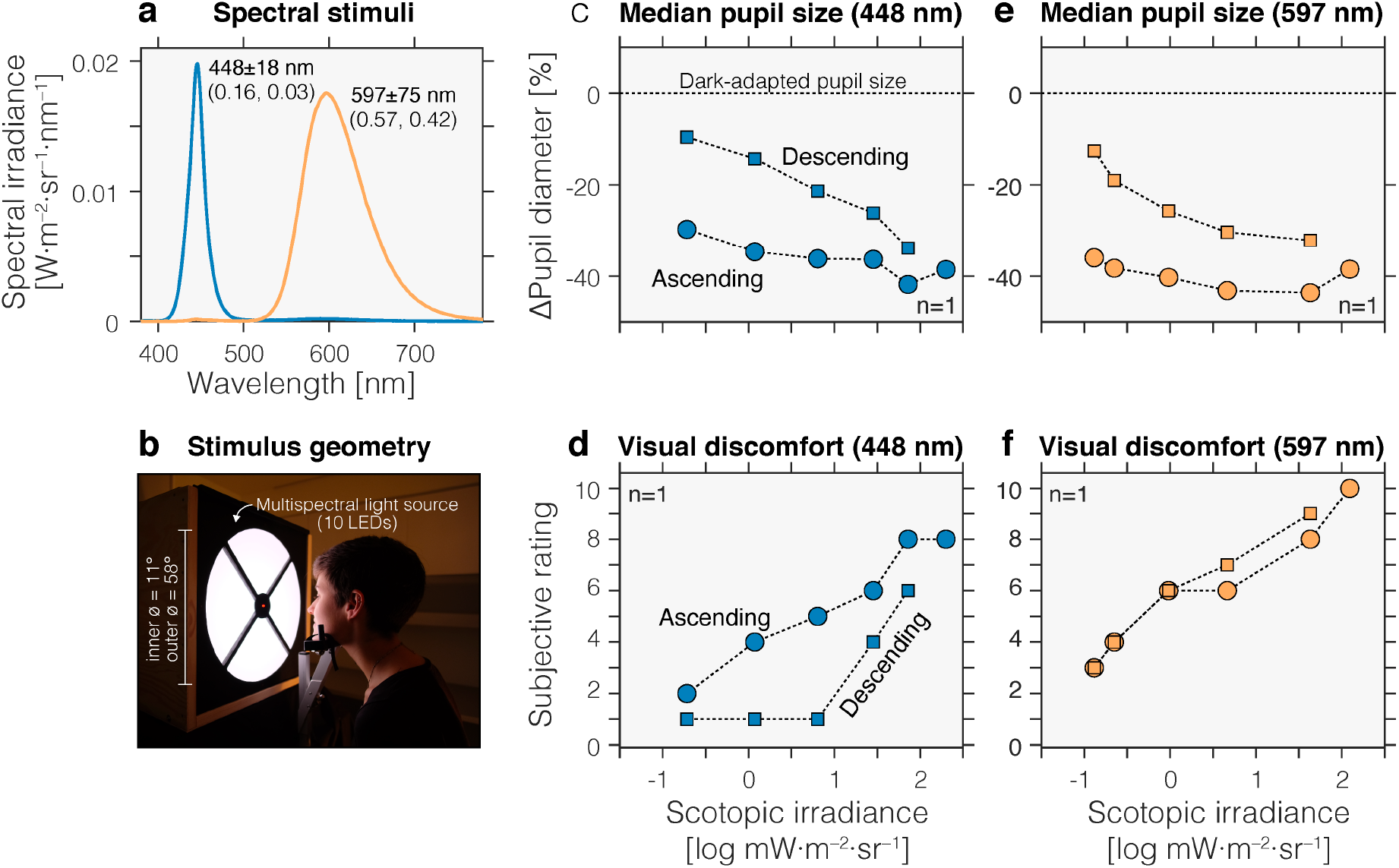
Visual discomfort and pupil responses in a congenital achromat (n=1). ***a*** Irradiance spectra of stimuli used, corresponding to a blue appearing and orange appearing light (to a trichromat). ***b*** Overview of stimulus geometry. Light emitted from a 10-primary tuneable LED-based light source was back-projected on a plexiglass surface, which the participant viewed in free-viewing conditions. ***c*** Pupil responses (relative to dark-adapted pupil size) in response to a series of ascending 448 nm stimuli, increasing in irradiance (squares), or to a series of descending light stimuli, decreasing in irradiance (circles). ***d*** Visual discomfort ratings in response to a series of ascending 448 nm stimuli, increasing in irradiance (squares), or to a series of descending light stimuli, decreasing in irradiance (circles). ***e*** Pupil responses (relative to dark-adapted pupil size) in response to a series of ascending 597 nm stimuli, increasing in irradiance (squares), or to a series of descending light stimuli, decreasing in irradiance (circles). ***f*** Visual discomfort ratings in response to a series of ascending 597 nm stimuli, increasing in irradiance (squares), or to a series of descending light stimuli, decreasing in irradiance (circles).

**Supplementary Figure S4:**
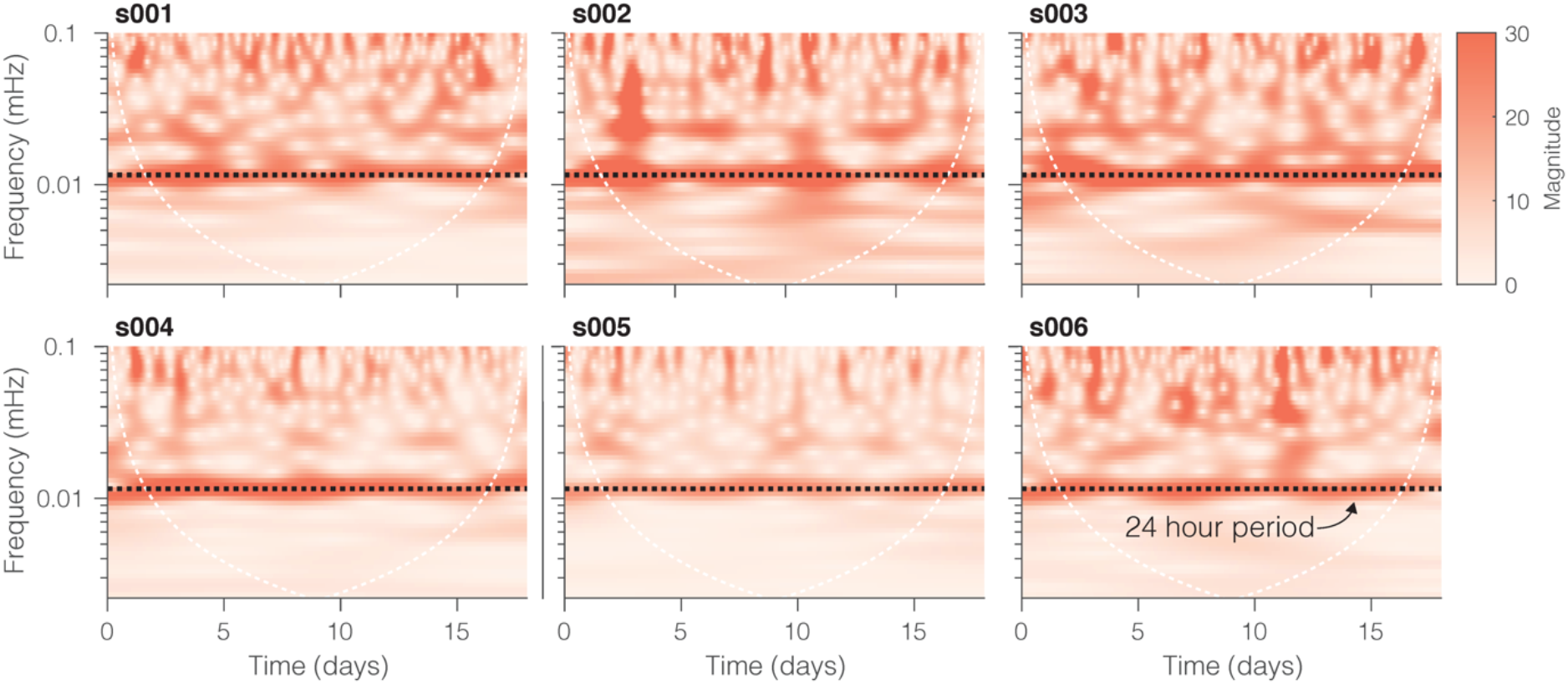
Analysis of actigraphy data accounting for non-stationarity confirms rest-activity cycles with 24-hour period. To understand the 24-hour periodicity of the measurements in the presence of possible non-stationarities (i.e. changes of phase and period throughout the protocol), actigraphy from all participants (n=6) were analysed using wavelets (25, 26). Activity data (PIMn) were subjected to the continuous wavelet transform using the analytic Morse wavelet with parameters γ = 3 (symmetry) and *p*^2^ = 60 (time-bandwidth product) implemented in MATLAB’s cwt function (55, 56). The black dashed line corresponds to the frequency associated to a 24-hour period. Any points outside of the white ‘U’-shaped curve are susceptible to edge effects.

**Supplementary Figure S5:**
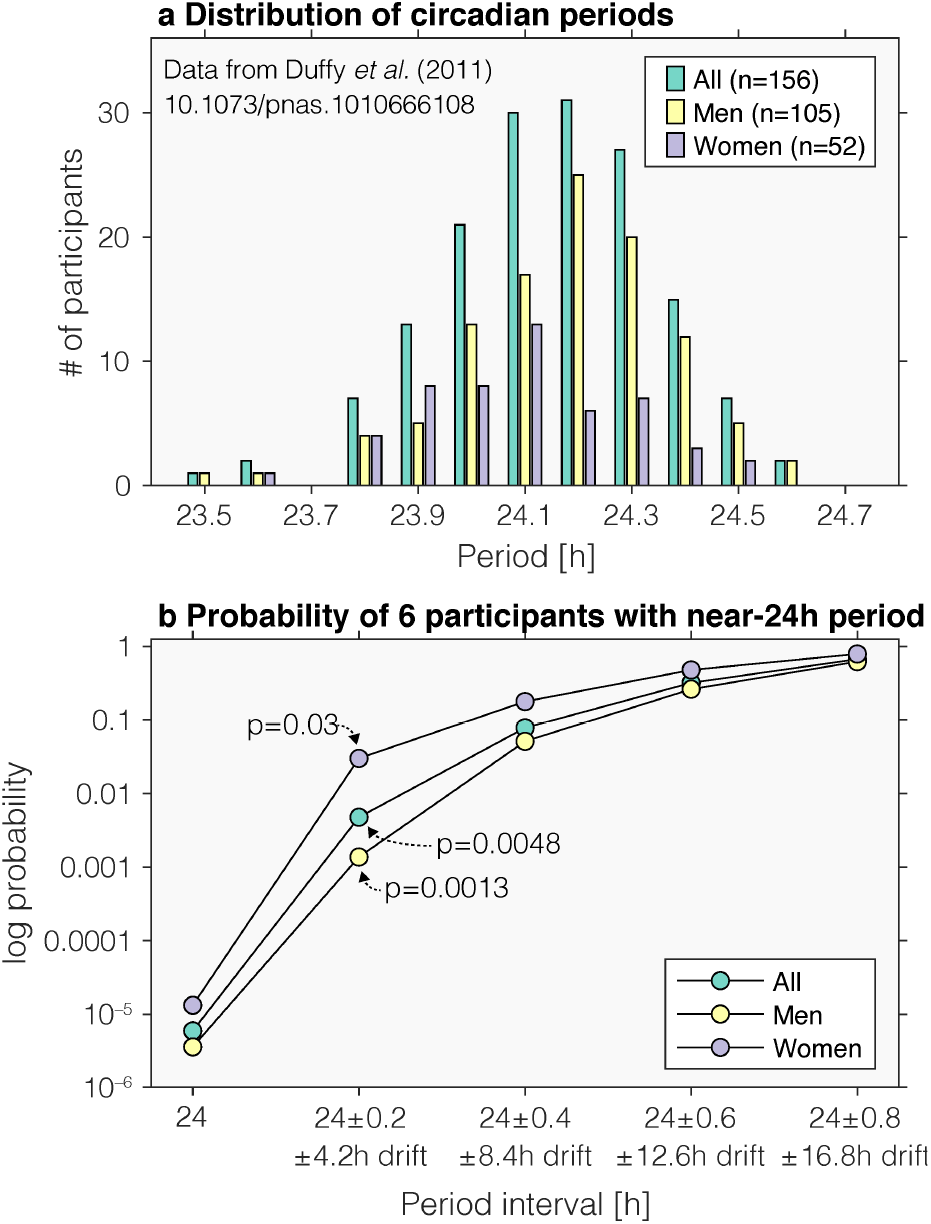
Probability of free-running participants with precise or near-24h period. ***a*** Distribution of circadian periods derived from temperature measurements in a general population as reported by Duffy *et al.* (2011). Data are replotted from their Figures 1 (all data; top panel), and Figure 2 (data aggregated by sex). ***b*** Probability to have recruited a total of 6 participants with varying period intervals leading to a range of drift values across an observation epoch of 21 days. The probability to recruit a participant with a period within the 24h “bin” is <0.0001, independent of whether data all data are considered, or disaggregated by sex. Assuming a more liberal criterion, the probability of recruiting a participant with the 24±0.2h bin (leading to a drift of ±4.2h over 21 days) is of course higher, and given by 0.1346^*6*^=0.0048 for six participants regardless of sex [0.1538^*n*^=0.03 for an all-female participant sample of size six, and 0.1238^*n*^=0.0013 for an all-male sample of size six (arrows indicated in figure)]. We validated these numbers numerically using repeated sampling from the empirical histogram using the inverse transform method and calculating the fraction that all six participants were landed in the period bin 24±0.2h (k=10,000 draws of six participants, m=30 repeats). The probability estimates for sampling six participants in the 24±0.2h bin were 0.0051±0.0007 (both male and female), 0.0301±0.0017 (all female), and 0.0014±0.0004 (all male), respectively. In this experiment, where we had five female and one male participant, we estimate the probability to randomly sample participants with a period between 23.9 and 24.1h to be 0.1538*5×*0.3333 = 0.018.

## References

1. J. F. Duffy, C. A. Czeisler, Effect of Light on Human Circadian Physiology. Sleep Med Clin 4, 165–177 (2009).

2. J. F. Duffy, K. P. Wright, Jr., Entrainment of the human circadian system by light. J Biol Rhythms 20, 326–338 (2005).

3. M. E. Jewett, D. B. Forger, R. E. Kronauer, Revised limit cycle oscillator model of human circadian pacemaker. J Biol Rhythms 14, 493–499 (1999).

4. M. Spitschan, Melanopsin contributions to non-visual and visual function. Curr Opin Behav Sci 30, 67–72 (2019).

5. A. Stockman, L. T. Sharpe, Into the twilight zone: the complexities of mesopic vision and luminous efficiency. Ophthalmic Physiol Opt 26, 225–239 (2006).

6. S. Kohl, C. Hamel, Clinical utility gene card for: Achromatopsia - update 2013. Eur J Hum Genet 21(2013).

7. M. H. Remmer, N. Rastogi, M. P. Ranka, E. J. Ceisler, Achromatopsia: a review. Curr Opin Ophthalmol 26, 333–340 (2015).

8. R. F. Hess, L. T. Sharpe, K. Nordby, Night vision : basic, clinical, and applied aspects (Cambridge University Press, Cambridge; New York, 1990), pp. xii, 550 p.

9. J. Aboshiha et al., A Quantitative and Qualitative Exploration of Photoaversion in Achromatopsia. Invest Ophthalmol Vis Sci 58, 3537–3546 (2017).

10. M. C. Aguilar et al., Automated instrument designed to determine visual photosensitivity thresholds. Biomed Opt Express 9, 5583–5596 (2018).

11. G. Schwerdtfeger, M. Gräf, Kantenfilterkontaktlinse und Kantenfiltergläser bei Achromatopsie. Z Prakt Augenheilkd 15, 322–328 (1994).

12. N. Angier (1992) New Clue to Vision: People Whose Glasses Must Be Rose-Colored. in New York Times.

13. F. Futterman, Understanding and coping with achromatopsia. (2004).

14. C. A. Czeisler et al., Suppression of melatonin secretion in some blind patients by exposure to bright light. N Engl J Med 332, 6–11 (1995).

15. J. T. Hull, C. A. Czeisler, S. W. Lockley, Suppression of Melatonin Secretion in Totally Visually Blind People by Ocular Exposure to White Light: Clinical Characteristics. Ophthalmology 125, 1160–1171 (2018).

16. J. Charng et al., Pupillary Light Reflexes in Severe Photoreceptor Blindness Isolate the Melanopic Component of Intrinsically Photosensitive Retinal Ganglion Cells. Invest Ophthalmol Vis Sci 58, 3215–3224 (2017).

17. J. Margraf, J. C. Cwik, V. Pflug, S. Schneider, Strukturierte klinische Interviews zur Erfassung psychischer Störungen über die Lebensspanne. Zeitschrift für Klinische Psychologie und Psychotherapie 46, 176–186 (2017).

18. K. V. Danilenko, E. G. Verevkin, V. S. Antyufeev, A. Wirz-Justice, C. Cajochen, The hockey-stick method to estimate evening dim light melatonin onset (DLMO) in humans. Chronobiol Int 31, 349–355 (2014).

19. D. J. Buysse, C. F. Reynolds, 3rd, T. H. Monk, S. R. Berman, D. J. Kupfer, The Pittsburgh Sleep Quality Index: a new instrument for psychiatric practice and research. Psychiatry Res 28, 193–213 (1989).

20. M. W. Johns, A new method for measuring daytime sleepiness: the Epworth sleepiness scale. Sleep 14, 540–545 (1991).

21. A. Zavada, M. C. Gordijn, D. G. Beersma, S. Daan, T. Roenneberg, Comparison of the Munich Chronotype Questionnaire with the Horne-Ostberg’s Morningness-Eveningness Score. Chronobiol Int 22, 267–278 (2005).

22. J. A. Horne, O. Ostberg, A self-assessment questionnaire to determine morningness-eveningness in human circadian rhythms. Int J Chronobiol 4, 97–110 (1976).

23. C. M. Mangione et al., Development of the 25-item National Eye Institute Visual Function Questionnaire. Arch Ophthalmol 119, 1050–1058 (2001).

24. J. D. Verriotto et al., New Methods for Quantification of Visual Photosensitivity Threshold and Symptoms. Transl Vis Sci Technol 6, 18(2017).

25. T. L. Leise, M. E. Harrington, Wavelet-based time series analysis of circadian rhythms. J Biol Rhythms 26, 454–463 (2011).

26. T. L. Leise, Wavelet analysis of circadian and ultradian behavioral rhythms. J Circadian Rhythms 11, 5(2013).

27. E. J. Van Someren et al., Bright light therapy: improved sensitivity to its effects on rest-activity rhythms in Alzheimer patients by application of nonparametric methods. Chronobiol Int 16, 505–518 (1999).

28. G. Hammad, M. Reyt (2020) pyActigraphy (feature/actrust branch, commit 598ef80). (https://github.com/ghammad/pyActigraphy).

29. K. Plum, Kantenfilter und seitlicher Blendschutz – ein praktischer Ratgeber (Gemeinsame Empfehlungen der Pro Retina und der WVAO) (Pro Retina Deutschland e.V., Bonn, 2019).

30. G. Bundesauschuss’ (2020) Richtlinie des Gemeinsamen Bundesausschusses über die Verordnung von Hilfsmitteln in der vertragsärztlichen Versorgung (Hilfsmittel-Richtlinie/HilfsM-RL). in BAnz AT 14.02.2020 B2, ed B. d. J. u. f. Verbraucherschutz (Bundesministerium der Justiz und für Verbraucherschutz, Berlin).

31. T. Roenneberg, L. K. Pilz, G. Zerbini, E. C. Winnebeck, Chronotype and Social Jetlag: A (Self-) Critical Review. Biology (Basel) 8(2019).

32. A. I. Luik, L. A. Zuurbier, A. Hofman, E. J. Van Someren, H. Tiemeier, Stability and fragmentation of the activity rhythm across the sleep-wake cycle: the importance of age, lifestyle, and mental health. Chronobiol Int 30, 1223–1230 (2013).

33. V. Bromundt et al., Sleep-wake cycles and cognitive functioning in schizophrenia. Br J Psychiatry 198, 269–276 (2011).

34. M. Wittmann, J. Dinich, M. Merrow, T. Roenneberg, Social jetlag: misalignment of biological and social time. Chronobiol Int 23, 497–509 (2006).

35. H. J. Burgess, J. K. Wyatt, M. Park, L. F. Fogg, Home Circadian Phase Assessments with Measures of Compliance Yield Accurate Dim Light Melatonin Onsets. Sleep 38, 889–897 (2015).

36. C. I. Eastman, V. A. Tomaka, S. J. Crowley, Circadian rhythms of European and African-Americans after a large delay of sleep as in jet lag and night work. Sci Rep 6, 36716(2016).

37. K. Brismar, B. Hylander, K. Eliasson, S. Rossner, L. Wetterberg, Melatonin secretion related to side-effects of beta-blockers from the central nervous system. Acta Med Scand 223, 525–530 (1988).

38. K. P. Wright, Jr., R. J. Hughes, R. E. Kronauer, D. J. Dijk, C. A. Czeisler, Intrinsic near-24-h pacemaker period determines limits of circadian entrainment to a weak synchronizer in humans. Proc Natl Acad Sci U S A 98, 14027–14032 (2001).

39. G. V. Vartanian et al., Melatonin Suppression by Light in Humans Is More Sensitive Than Previously Reported. J Biol Rhythms 30, 351–354 (2015).

40. N. N. Takasu et al., Repeated exposures to daytime bright light increase nocturnal melatonin rise and maintain circadian phase in young subjects under fixed sleep schedule. Am J Physiol Regul Integr Comp Physiol 291, R1799–1807 (2006).

41. C. Cajochen, M. E. Jewett, D. J. Dijk, Human circadian melatonin rhythm phase delay during a fixed sleep-wake schedule interspersed with nights of sleep deprivation. J Pineal Res 35, 149–157 (2003).

42. T. Roenneberg, T. Kantermann, M. Juda, C. Vetter, K. V. Allebrandt, “Light and the human circadian clock” in Handb Exp Pharmacol, A. Kramer, M. Merrow, Eds. (2013), 10.1007/978-3-642-25950-0_13, pp. 311–331.

43. T. Roenneberg, C. J. Kumar, M. Merrow, The human circadian clock entrains to sun time. Curr Biol 17, R44–45 (2007).

44. F. L. Ruberg et al., Melatonin regulation in humans with color vision deficiencies. J Clin Endocrinol Metab 81, 2980–2985 (1996).

45. H. Calligaro et al., Rods contribute to the light-induced phase shift of the retinal clock in mammals. PLoS Biol 17, e2006211 (2019).

46. M. C. Gimenez, D. G. Beersma, P. Bollen, M. L. van der Linden, M. C. Gordijn, Effects of a chronic reduction of short-wavelength light input on melatonin and sleep patterns in humans: evidence for adaptation. Chronobiol Int 31, 690–697 (2014).

47. R. P. Najjar et al., Aging of non-visual spectral sensitivity to light in humans: compensatory mechanisms? PLoS One 9, e85837 (2014).

48. J. F. Duffy, R. E. Kronauer, C. A. Czeisler, Phase-shifting human circadian rhythms: influence of sleep timing, social contact and light exposure. J Physiol 495 (Pt 1), 289–297 (1996).

49. R. E. Mistlberger, D. J. Skene, Nonphotic entrainment in humans? J Biol Rhythms 20, 339–352 (2005).

50. S. M. T. Wehrens et al., Meal Timing Regulates the Human Circadian System. Curr Biol 27, 1768–1775 e1763 (2017).

51. K. Krauchi, C. Cajochen, E. Werth, A. Wirz-Justice, Alteration of internal circadian phase relationships after morning versus evening carbohydrate-rich meals in humans. J Biol Rhythms 17, 364–376 (2002).

52. CIE (2018) CIE S 026/E:2018: CIE System for Metrology of Optical Radiation for ipRGC-Influenced Responses to Light. (CIE Central Bureau, Vienna, Austria).

53. K. W. Houser, M. Wei, A. David, M. R. Krames, X. S. Shen, Review of measures for light-source color rendition and considerations for a two-measure system for characterizing color rendition. Opt Express 21, 10393–10411 (2013).

54. ISO, ISO 23539:2005(E)/CIE S 010/E:2004: Photometry —The CIE system of physical photometry (2005).

55. S. C. Olhede, A. T. Walden, Generalized Morse wavelets. IEEE Transactions on Signal Processing 50, 2661–2670 (2002).

56. J. M. Lilly, S. C. Olhede, Higher-Order Properties of Analytic Wavelets. IEEE Transactions on Signal Processing 57, 146–160 (2009).

